# Treating Porcine Abscesses With Histotripsy: A Pilot Study

**DOI:** 10.1101/2020.09.25.314302

**Authors:** Thomas J. Matula, Yak-Nam Wang, Tatiana Khokhlova, Daniel F. Leotta, John Kucewicz, Andrew A. Brayman, Matthew Bruce, Adam D. Maxwell, Brian E. MacConaghy, Gilles Thomas, Valery P. Chernikov, Sergey V. Buravkov, Vera A. Khokhlova, Keith Richmond, Keith Chan, Wayne Monsky

**Affiliations:** Applied Physics Laboratory, University of Washington, Seattle, WA 98105, USA; Department of Gastroenterology, University of Washington, Seattle, WA 98105, USA; Department of Urology, University of Washington, Seattle, WA 98105, USA; Research Institute of Human Morphology, Laboratory of Cell Pathology, Moscow, Russia; Faculty of Fundamental Medicine, M.V. Lomonosov Moscow State University, Moscow, Russia; Department of Acoustics, Physics Faculty, M.V. Lomonosov Moscow State University, Moscow Russia; BioDevelopment Associates, Stanwood, WA, 98043, USA; Department of Radiology, University of Washington, Seattle, WA 98105, USA

**Author notes:** Corresponding Author: Thomas J. Matula, Applied Physics Laboratory, 1013 N.E. 40th Street, Seattle WA 98105, USA.

**Keywords:** Histotripsy, Abscess, Focused Ultrasound, Image-Guided Therapy, HIFU, Acoustic bactericide, Bacteria inactivation, Focused Ultrasound

## Abstract

Infected abscesses are walled-off collections of pus and bacteria. They are a common *sequela* of complications in the setting of surgery, trauma, systemic infections, and other disease states. Current treatment is typically limited to antibiotics with long-term catheter drainage, or surgical wash-out when inaccessible to percutaneous drainage or unresponsive to initial care efforts. Antibiotic resistance is also a growing concern. Although bacteria can develop drug resistance, they remain susceptible to thermal and mechanical damage. In particular, short pulses of focused ultrasound (*i*.*e*., histotripsy) generate mechanical damage through localized cavitation, representing a potential new paradigm for treating abscesses non-invasively, without the need for long-term catheterization and antibiotics. In this pilot study, boiling and cavitation histotripsy treatments were applied to subcutaneous and intramuscular abscesses developed in a novel porcine model. Ultrasound imaging was used to evaluate abscess maturity, for treatment monitoring and assessment of post-treatment outcomes. Disinfection was quantified by counting bacteria colonies from samples aspirated before and after treatment. Histopathological evaluation of the abscesses was performed to identify changes resulting from histotripsy treatment and potential collateral damage. Cavitation histotripsy was more successful in reducing the bacterial load while having a smaller treatment volume compared with boiling histotripsy. The results of this pilot study suggest focused ultrasound may lead to a technology for *in situ* treatment of acoustically accessible abscesses.

## Introduction

The Agency for Healthcare Research and Quality reported that in 2011 alone, >262,000 U.S. patients were hospitalized for abscess treatment, accruing >$6.2B in aggregate U.S. hospital charges (Chan 2013). Abscesses may arise from diverticulitis, appendicitis (Park and Charles 2012, Singh and Khardori 2012), gastric bypass surgery (Sarkhosh, et al. 2013), hepatic resection (Vladov, et al. 2013, Zimmitti, et al. 2013), chemoembolization (Woo, et al. 2013), radio frequency ablation (Taner, et al. 2013), pancreatectomy (Yui, et al. 2013), otitis media post-surgical complications (Yorgancilar, et al. 2013), and colorectal surgery (Li, et al. 2013), to name a few causes. Abscesses can occur anywhere, including the lung (Seo, et al. 2013) or mediastinum (Leong, et al. 2013), brain (Radoi, et al. 2013, Saez-Llorens and Guevara 2013), prostate (Roux, et al. 2013), spinal cord (Urrutia, et al. 2013), perianal tissues (Morcos, et al. 2013), and skin and soft tissue (Brook 2008, Cardona and Wilson 2015). Abdominal abscesses rank among the top 10 diseases for the highest 30-day rehospitalization rate (Elixhauser and Steiner 2013).

Abscess care depends on size, location, and complexity, among other patient factors. Superficial abscesses are most often treated by incision and drainage, even though it is painful and scarring (Mistry 2013). For abdominal abscesses, since the early 1980s, percutaneous catheter drainage replaced open surgery following the pioneering reports of Haaga *et a*l (Haaga, et al. 1977, Haaga and Weinstein 1980) and Gerzof *et al*, (Gerzof, et al. 1979, Gerzof, et al. 1981) and is today the current standard of care (Rivera-Sanfeliz 2008). Percutaneous catheter drainage typically involves inserting a drainage catheter into the abscess under CT or ultrasound guidance; it usually remains in place for up to several weeks, during which repeat drainage, drain manipulation (including upsizing of drains), follow-up CT or sinogram may be needed. In addition, percutaneous drainage is often unsuccessful when the contents are too viscous (Lorenz and Thomas 2006). Quality of life can be significantly affected adversely with the placement of any drain (Monsky, et al. 2014). Patients experience significant pain and discomfort, and even simple daily activities like sitting and walking can be difficult. Complications such as clogged drains, secondary infections or having the drain jar loose require rehospitalization and wound management. Abscesses which are difficult to drain due to loculation, viscosity, close proximity to other vital structures or without a safe window for percutaneous drainage may require more invasive surgical washout procedures.

Focused ultrasound has the potential to treat some of these infected fluid collections with an associated reduction in related morbidity and complications. Intense cavitation homogenizes cells, leaving liquefied cellular debris (Maxwell 2012, Wang, et al. 2013, Khokhlova, et al. 2014, Khokhlova, et al. 2016, Wang, et al. 2018). Also, macroscopic streaming circulates cells and other particles through the focus, effectively increasing the treatment volume.

Acoustic inactivation of bacteria has been a topic of interest in a variety of fields ranging from food preservation to medicine for over 90 years (Harvey and Loomis 1929). An overview of the primary publications and review articles dealing with acoustic inactivation of bacteria is described by Brayman, et. al. (Brayman, et al. 2017). Bacterial inactivation by intense (non-thermal) ultrasound - termed histotripsy - results from inertial cavitation *via* direct mechanical effects related to the shear forces associated with bubble collapse (Gao, et al. 2014) and *via* sonochemical effects (Joyce, et al. 2003). Earlier results obtained with *E. coli* in suspension are consistent with this general conclusion (Brayman, et al. 2017, Brayman, et al. 2018). Generally, inactivation rates increased above a pressure threshold (P^−^ ∼ 7 MPa) and with exposure time. TEM images of treated bacteria (*E. coli*) showed mechanical shearing of the membrane, disruption of the cytoplasm, and cellular debris.

Much of the focused ultrasound research has been directed to treat biofilms (Bigelow, et al. 2009, Bigelow, et al. 2017) and solid tumors (Orsi, et al. 2010, Al-Bataineh, et al. 2012). To ablate tumors, not only does the temperature need to be elevated (and controlled), but the focal spot must be steered to cover the entire tumor, extending into the tumor margins. Conversely, heating is more difficult in liquids. Instead, histotripsy generates cavitation that mechanically homogenizes tissue. Streaming induced at the focal area via the acoustic radiation force brings new material into the focus (Maxwell, et al. 2009). Cavitation greatly enhances streaming due to the large interaction between the sound wave and bubbles. General streaming patterns can lead to liquefaction of the viscous contents. Bubble oscillations themselves create microstreaming patterns that are known to disrupt cells due to high transient shear forces (Wu and Nyborg 2008). Thus, streaming and microstreaming significantly increases the cavitation activation zone. Unlike the procedure for tumors, histotripsy treatment for abscesses should not require that the focus be moved near the abscess capsule. This was demonstrated in an *in vitro* study which used a fixed-focus system to treat a 10-cc sample by drawing bacteria into the comparatively small focal volume (Brayman, et al. 2018).

Because histotripsy has never been applied to relatively large viscous liquid pus abscesses *in situ*, these preliminary studies focused on evaluating targeting and assessing treatment efficacy from cavitation histotripsy (CH) or Boiling Histotripsy (BH) using a novel porcine model (Wang, et al. 2020). Both *ex vivo* and *in vivo* treatments of subcutaneous and intramuscular abscesses created in the porcine model are presented. The abscesses were only partially treated so that ultrasound imaging could be used to assess gross changes in the treatment zone. Target evaluation was performed by quantifying cavitation zones and histological assessments. Treatment efficacy was quantified by collecting and counting bacteria colonies before or after treatment. Several representative *ex vivo* and *in vivo* cases are described herein.

## Materials and Methods

### Animal Model

All procedures were approved by the Institutional Animal Care and Use Committee (IACUC) at R&R Rabbitry (Stanwood, WA). Female domestic swine (Yorkshire/Hampshire cross, age 3-6 months, weight 45-65 kg) were used in this study. In a recent paper (Wang, et al. 2020), we described the development of a porcine animal model in which multiple large multiloculated bimicrobial abscesses can be formed at distinct sites in the same animal. These abscesses satisfy the true definition of an abscess (Schein, et al. 1997, Kumar, et al. 2015) and are formed within a short period. Briefly, on the day of inoculation, the bacterial mix, consisting of the facultative anaerobe *E. coli* (strain ATCC 25922, ATCC, Manassas VA, USA), the obligate anaerobe *B. fragilis* (strain ATCC 23745) and sterile dextran micro carrier bead (Cytodex®, Millipore Sigma, St. Louis, MO) suspension in a ratio of 1:1:2 was prepared so that the final inoculum contained 10^6^ cfu/ml *E. coli* and 10^6^ cfu/ml *B. fragilis* (cfu = colony forming units). Bacterial concentration was determined using spectrophotometric absorbance readings and confirmed with Compact Dry(tm) EC100 and TC assay plates (Hardy Diagnostics, Santa Maria, CA USA).

Animals were sedated (Telazol/Ketamine/Xylazine) and maintained under a surgical plane of anesthesia with isoflurane. All animals were instrumented to monitor heart rate, ECG, blood oxygen saturation and temperature. The haunches, abdomen and in some animals, the neck of the animal were depilated, cleaned and disinfected with consecutive rounds of 70% isopropanol. Injections were performed under ultrasound guidance into 6 to 8 separate sites in a single animal: 1.5 cm deep into the left and right biceps femoris, left and right brachiocephalicus (in one animal), and subcutaneously on either side of the midline. Before the intramuscular injections, the tip of the needle was moved back and forth fifty times in a fanning motion to create a small pocket of injury before injecting. For each animal 10 ml of the inoculate were injected at each site and the location was marked with dye. After the inoculations, the animals were recovered and monitored every day for signs of pain or distress, reduction in appetite, loss of weight and mobility. Abscesses were allowed to form over 3-6 weeks. Pigs that were recovered after abscess treatment showed no signs of fever, bacteremia, or sepsis.

### Ultrasound imaging of abscesses

Growth and development of abscesses were monitored at least weekly with ultrasound imaging. In addition, ultrasound imaging was used to assess the abscess at the time of treatment. Several ultrasound platforms were used throughout the experiments, depending on machine availability and the type of imaging performed: LOGIQ Book XP with 8L-RS 4-12 MHz linear array probe (GE Healthcare, Milwaukee, WI, USA), Aixplorer with L14-5 probe (Supersonic Imagine, Aix-en-Provence, France), or, for treatment monitoring, the Sonix RP with a P7-4 phased array probe (Ultrasonix Medical, Richmond, BC, Canada). With varying image quality, the same abscessal features could be successfully visualized with all platforms: a well-circumscribed, avascular, mixed echogenic core surrounded by a vascularized hypoechoic capsule. In some cases, palpation with the ultrasound probe was used to observe movement of the abscessal contents as a gross indication of the liquidity of the pus. Modalities used included B-mode (for general and 3D imaging and to monitor treatment), color and power Doppler (to evaluate vascularity), and Shearwave elastography (SWE, to evaluate stiffness). For *ex vivo* studies, 3D imaging was performed using a linear array probe (L12-5, HDI 5000, Philips Ultrasound, Bothell, WA, USA) integrated with a magnetic tracking system (Flock of Birds, Ascension Technology Corp., Burlington, VT, USA). Custom software recorded the 3D location and orientation of a series of 2D image planes acquired while scanning the abscess, and a 3D volume was reconstructed from the 2D images (Leotta, et al. 2018). This 3D imaging method was also performed for *in vivo* studies, with the Supersonic Imagine L14-5 probe tracked using a second magnetic tracking system (trakSTAR, NDI, Waterloo, ON, Canada).

## Experimental setup and procedures

### Ex vivo studies

*Ex vivo* treatments allowed for a simpler procedure to align the tissue sections with the treatment volume and enable more precise translation of the ultrasound focus throughout the volume. For these studies, the animals were euthanized 6 weeks post inoculation, and abscesses were dissected out *en bloc* with the overlying skin intact. The excised tissue was placed in degassed saline for treatment on the same day. The samples were embedded into 1.5% Weight/Volume low melting point agarose gel for ease of handling. Sampling of the abscessal contents was attempted with an 18G needle under ultrasound guidance. 3D ultrasound imaging was performed and the agarose embedded abscess was placed into a custom built holder attached to an automated 3D positioning system. Treatments were performed in a large tank filled with degassed deionized water, at room temperature. The integrated imaging-therapy probe shown in Fig. 1 as part of the in vivo setup was mounted in the water tank for these studies. The sample was oriented such that the skin surface was facing the treatment transducer and was perpendicular to the acoustic propagation axis. The histotripsy focus was shifted through the volume of the abscess in a 2D raster grid, in the plane perpendicular to the transducer axis. Immediately after histotripsy treatment, 3D ultrasound imaging of the abscess was performed again and a small sample of abscessal contents in the treatment region was aspirated with an 18-G needle under ultrasound guidance. Pre- and post-treatment samples were plated and colonies counted for comparative assessments of treatment efficacy.

**Figure 1.**
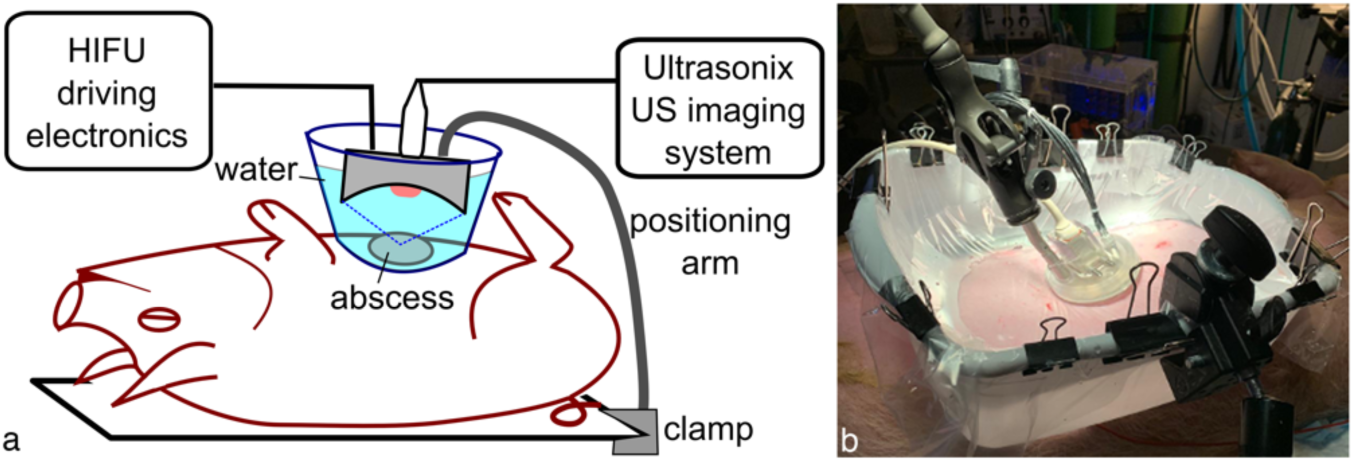
Illustration of the ultrasound-guided histotripsy setup for in vivo abscess treatment. For acoustic coupling to the skin, the histotripsy treatment head was immersed in a water container with acoustically transparent window positioned over the abscess. The treatment head was attached to either a flexible positioning (or robot) arm that allowed manual (or automated) movement of the transducer focus across the targeted abscess while delivering a specified treatment protocol at each focus location.

### In vivo studies

Histotripsy treatment *in vivo* was performed in 5 animals (10 mature abscesses). On the day of the treatment, the animal was sedated, intubated and maintained under a surgical plane of anesthesia with isofluorane. Pigs were placed in a supine or lateral recumbent position on the surgical table, depending on which abscess was treated. The area around the abscess(es) to be treated was prepared as described above. Where possible, ultrasound imaging was performed to assess structural features, 3D volume, vascularity and stiffness. Specific abscesses to be treated were selected based on the aforementioned evaluation. Before each treatment, attempts were made to aspirate small abscessal samples (<50 µL) with an 18G needle under ultrasound guidance. This procedure was repeated after treatment, and the pre- and post-treatment samples were plated and colonies counted for comparative assessments of treatment efficacy.

The histotripsy treatment setup for *in vivo* studies is illustrated in Fig. 1. A bath consisting of a plastic coupling container with the bottom removed was lined with a 38-µm thick low-density polyethylene membrane. The skin was wetted with degassed water containing 1% isopropyl alcohol, and a thin layer of ultrasound gel that had been centrifuged to remove large air bubbles was applied to the skin. Degassed water was used to wet the surface of the gel layer before the coupling container was put in place. The container was then filled with degassed water and any visible air bubbles in the gel between the membrane and skin were removed. All treatments were performed with one of three custom-built HIFU transducers with a circular central opening that incorporated a coaxially aligned ultrasound imaging probe for targeting and in-treatment B-mode cavitation monitoring. The parameters of the transducers and the corresponding imaging probes are summarized in Table 1; one of two imaging systems was used for in-treatment imaging - GE LOGIQ Book XP or Sonix RP. The HIFU treatment head was mounted on a flexible positioning arm that could be locked in place for histotripsy exposure, and then unlocked again to manually move between treatment locations. In some treatments the manually manipulated positioning arm was replaced by a collaborative robotic arm (UR3e, Universal Robots GmbH, Odense, Denmark), such that the treatment locations could be more precisely controlled. To deliver histotripsy exposures, the transducers were driven with custom high power electronics built in house as described in detail elsewhere (Maxwell, et al. 2017).

**Table 1.**
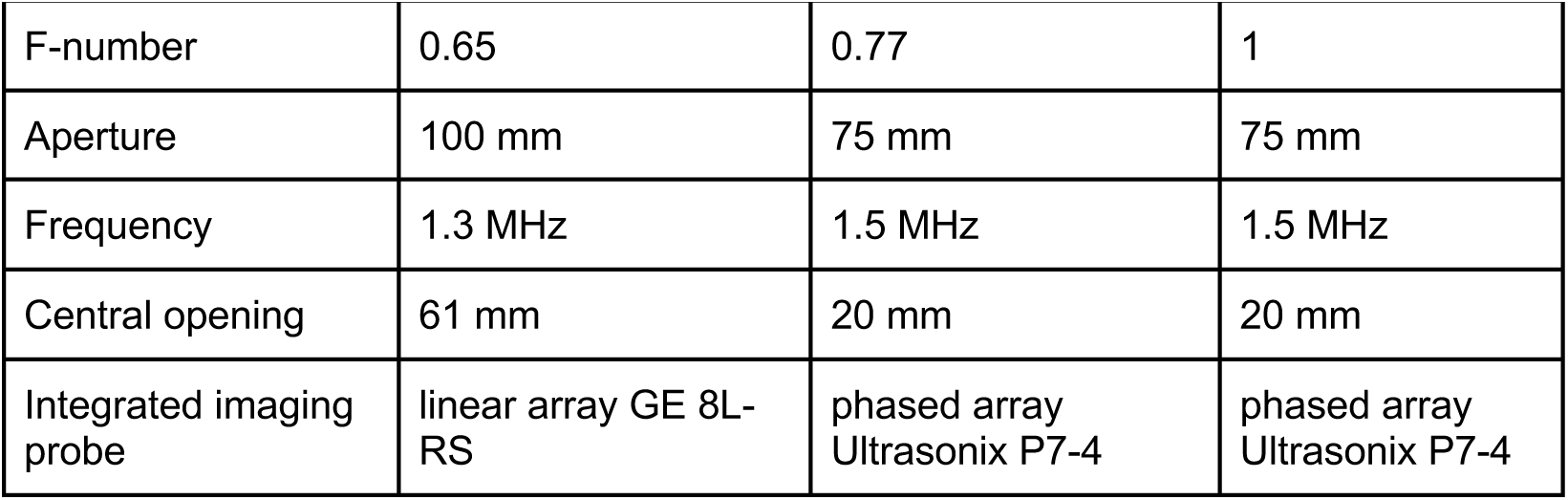
Parameters of the three histotripsy transducers used in the study. All transducers were 12-element sector arrays with a circular central opening built in house.

|  |  |  |  |
| --- | --- | --- | --- |
| F-number | 0.65 | 0.77 | 1 |
| Aperture | 100 mm | 75 mm | 75 mm |
| Frequency | 1.3 MHz | 1.5 MHz | 1.5 MHz |
| Central opening | 61 mm | 20 mm | 20 mm |
| Integrated imaging probe | linear array GE 8L-RS | phased array Ultrasonix P7-4 | phased array Ultrasonix P7-4 |

Immediately following histotripsy treatment, ultrasound imaging was performed to visualize any changes to the abscess produced by the treatment. After the animal was euthanized, treated abscesses were carefully removed *en bloc* and fixed in 10% neutral buffered formalin for histological evaluation.

### Histotripsy treatment protocols

Two types of histotripsy treatment protocols were used in the experiments: shock-scattering histotripsy (for brevity, we will refer to this regime as cavitation histotripsy, CH) and boiling histotripsy (BH). Both approaches use short bursts of high amplitude nonlinear acoustic waves delivered at low duty cycle to induce gas and/or vapor bubble activity at the focus that mechanically disintegrates tissue at the focus. CH uses microseconds-long pulses delivered at pulse repetition frequencies (PRFs) in the kilohertz range, with peak rarefactional focal pressures sufficient for cavitation nucleation, and high amplitude shock fronts forming at the focus (Maxwell, et al. 2011). In BH milliseconds-long acoustic pulses with high amplitude shocks forming at the focus are used to induce the formation of vapor bubble at the focus within each pulse. BH pulses are delivered at lower PRF, resulting in a duty cycle of 1-2% (Khokhlova, et al. 2011). Here, we have used different sets of CH and BH pulsing parameters for each abscess that fit within the constraints described above; they are summarized in Table 2.

**Table 2.** Pulsing protocols used in histotripsy treatments of abscesses

|  | Pulse duration | Duty cycle | Time per focus | Focus spacing |
| --- | --- | --- | --- | --- |
| CH | 7 or 33 μs | 1% | 2-5 min | 3-5 mm |
| BH | 7.7 or 10 ms | 1% | 1-3 min | 3-5 mm |

Both BH and CH have been extensively studied previously for soft tissue ablation, and the *in situ* pressures needed to achieve either regime are known. However, because the histotripsy target in the present case was abscess pus - a semi-liquid with likely elevated and unique dissolved gas content - the pulse-average acoustic power thresholds for either BH or CH were determined for each corresponding abscess prior to treatment. Specifically, a single focus location (the first location in the treatment grid) was sonicated with BH or CH pulses at gradually increasing amplitude until a hyperechoic region was observed at the focus, indicating boiling or cavitation cloud activity, respectively. The treatments that followed were performed slightly above that threshold (10–15% increase in driving voltage), with all points in the treatment grid receiving the same power, similarly to our previous studies (Khokhlova, et al. 2019). To estimate the corresponding *in situ* focal pressure waveforms, the waveforms measured in water for the three transducers by the fiber optic probe hydrophone (FOPH2000, RP Acoustics, Leutenbach, Germany) were derated to the corresponding depth (1-3 cm) in tissue with attenuation of 1.7 dB/cm using a modified derating approach for nonlinearly distorted waveforms as previously described (Bessonova, et al. 2010). Figure 2 shows representative derated focal waveforms for each of the transducers when used in both CH and BH regime (note: the transducer with F-number of 1 was only used in BH treatments). All waveforms contained a high-amplitude shock front, although peak compressional and rarefactional pressures were noticeably different for the different transducers. The *in situ* focal pressures required for BH and CH treatments were similar in abscess pus (p^+^=96 MPa, p^-^=18 MPa, shock amplitude 121 MPa for the transducer with F-number of 0.77, and p^+^=121, p^-^=20 MPa, shock amplitude 131 MPa for the transducer with F-number of 0.65, see Fig. 2), unlike most soft tissues, in which CH threshold is typically higher. The derated peak positive and peak negative pressures were also consistent across abscesses and differed from the ones shown in Fig. 2 and stated above by under 10 MPa and 1 MPa, respectively.

**Figure 2.**
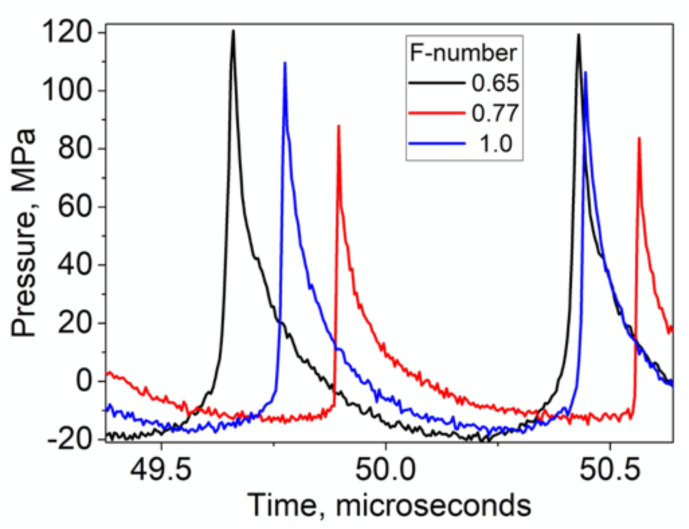
Representative in situ focal pressure waveforms used during histotripsy treatments (both CH and BH) by the three transducers. The waveforms were measured in water by FOPH and derated into corresponding depth in tissue. The in situ peak pressure levels were consistent across the treated abscesses. Note: the transducer with F-number of 1 was only used in BH exposures.

The treatment time per focus location also varied between different abscesses within 1-5 minutes and is provided in Table 2. While the spacing between locations could not be controlled precisely in the *in vivo* case, it was estimated through in-treatment B-mode imaging (see Table 2). That is, it was based on making the best effort to merge the areas of bubble activity in the adjacent location and provide a continuous treatment volume. The number of treatment locations depended on the size of the abscess and the treatment regime employed, ranging between 4-20.

### Histopathology

Fixed abscesses were grossed and processed to maintain the treatment zones as whole cross-sections of the abscesses as much as possible. Serial sections were stained with Hematoxylin and Eosin (H&E) for general tissue morphology, or Masson’s Trichrome stain to visualize the presence of a fibrous capsule Abscesses were evaluated for the presence of a connective tissue capsule; the extent of the capsule; damage to the capsule; the presence of cell debris and indication of histotripsy treatment.

### Bacteria viability

Abscessal contents that were sampled immediately before and after treatment were subjected to three serial dilutions using 3% Brain Heart Infusion broth using aliquots no smaller than 25 µL and a total dilution factor of 6.774 × 10^6^. One milliliter of each dilution was delivered to the surface of Compact DryTM EC100 plates (Hardy Diagnostics, Santa Maria, CA, USA) These plates specifically stain *E. coli* colonies deep blue, and other coliforms red. Colonies were counted to obtain a colony forming unit count per mL (cfu/mL). These counts were used to calculate changes in bacterial viability resulting from treatment.

### Transmission Electron Microscopy (TEM)

Three sample pairs (before and after treatment) were fixed in half-strength Karnovsky’s fixative for ultrastructural analysis. Methods similar to those used in Brayman et al. were used to process the samples (Brayman, et al. 2018). Briefly, samples were pelleted by centrifugation (8,000 g x 20 minutes). The pelleted samples were fixed with 1% osmium tetroxide before *en block* staining with 1% uranyl acetate and embedding in epon-araldite. Ultrathin sections were further staining with Reynolds lead citrate solution before being imaged on a transmission electron microscope (JEOL 101, Tokyo, Japan). Digital images were obtained with a Gatan camera.

## Results

### Abscess maturity

As the abscess matures from a region of inflammation to an encapsulated organized mass of viscous pus, it consolidates and a fibrous capsule is formed around the purulent material. Under B-mode ultrasound, the capsule appears more hypoechoic (*e*.*g*., Fig. 3a). The capsule is highly vascularized, as seen with color Doppler, while the variably echogenic core is devoid of vessels (Fig. 3b). These features were used to determine when the abscess was mature and organized and thus ready to treat. Not all abscesses grew large enough to be of clinical significance for treatment (>∼3 cm). Inoculating 6-8 regions consistently produced abscesses large and mature enough to treat. The maturation time was determined by quantifying the presence of a capsule. In most cases, the abscess matured in 3 weeks. Table 3 quantifies when an abscess was ready to treat for 21 different abscesses. Only 1 of the 21 abscesses failed to form a robust capsule that could be identified easily under B-mode or Doppler.

**Table 3.**
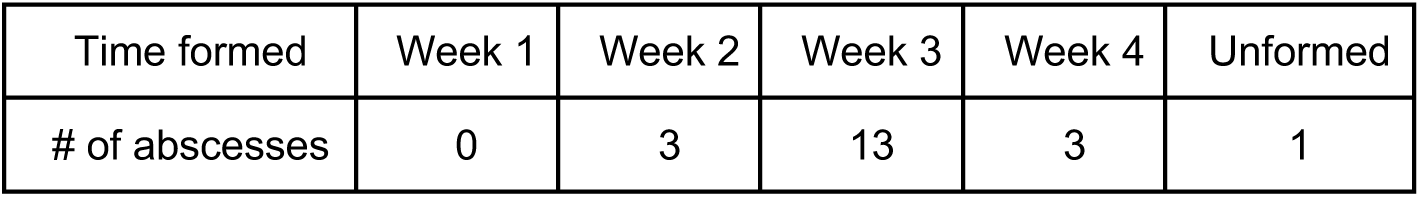
Capsule formation time, used to determine abscess maturity.

**Figure 3.**
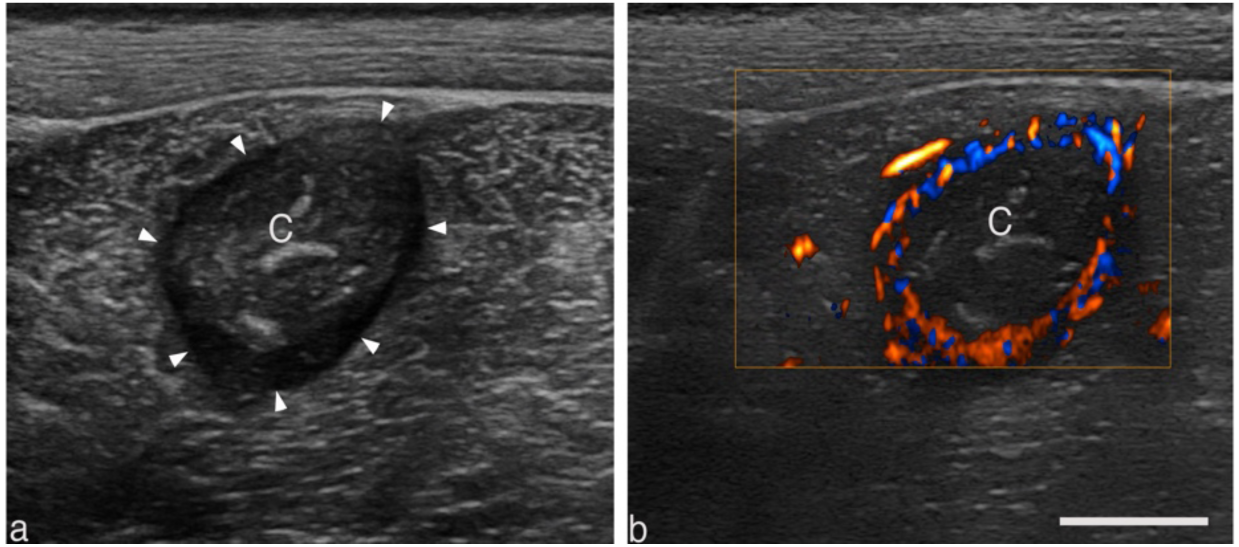
Representative B-mode (a) and color doppler (b) images of a well-circumscribed intramuscular abscess displaying salient features common to larger abscesses. The center of the abscess lies 2 cm below the skin surface within the biceps femoris. The variably echogenic core (C) is avascular and often surrounded by a vascular hypoechoic rim (arrowhead). Scale bar represents 1 cm.

### Ex vivo studies

The outcome of BH and CH treatments performed in *ex vivo* abscesses were visualized with post treatment ultrasound imaging and histological evaluation. An example of an abscess treated with both CH and BH is shown in Fig. 4. This abscess was formed by injection of the inoculate subcutaneously to the side of the midline of the pig. Before treatment, the abscess appeared as a distorted oblate spheroid in orthogonal image planes. Its dimensions were 22 × 36 × 37 mm. The abscessal contents appeared relatively uniform and more hypoechoic than the surrounding tissue.

**Figure 4.**
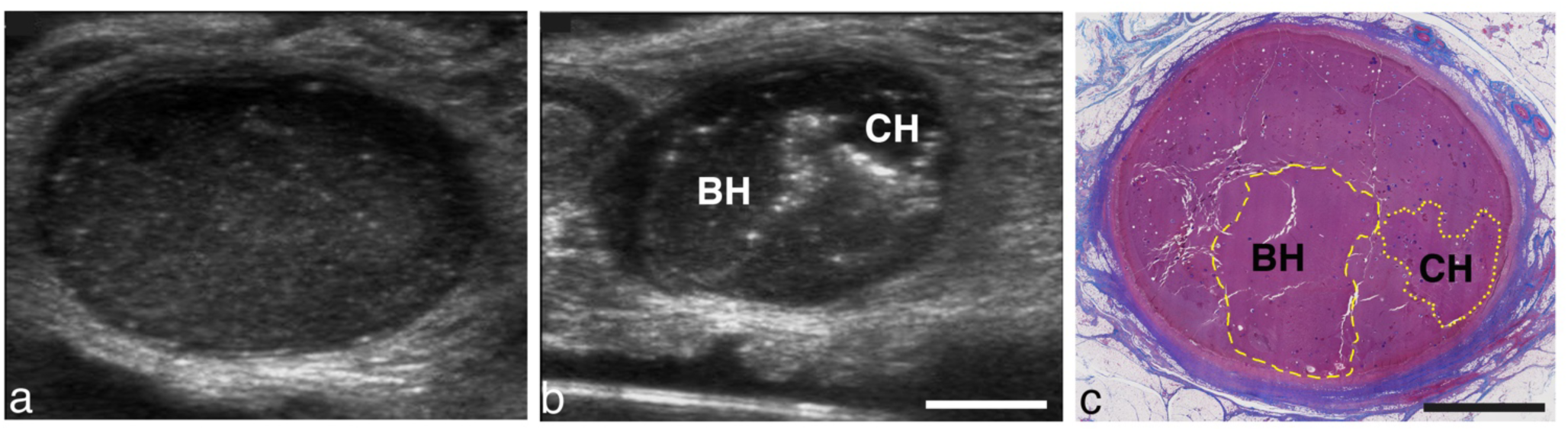
(a) Pre-treatment image of an abscess viewed along the long axis of the abscess. (b) Post treatment image viewed in a short-axis plane, orthogonal to the image plane in (a). Bubbles outline the focal regions from BH and CH treatments. (c) Masson’s trichrome stained section of the abscess in the short-axis plane. Collagen-rich connective tissue stains blue; nuclei stains black; muscle, cytoplasm and keratin pink-red. The abscess is surrounded by a collagen-rich capsule. The center of the abscess contains cells, tissue fragments and protein which appears pink at a low magnification. Yellow dashed or dotted line indicate the main border of the BH or CH lesion, respectively. Scale bar represents 1 cm.

BH treatment (10 ms bursts) was applied with the F#=0.77 transducer to 8 locations (2*x*4 rectangular grid) separated by 3 mm, for 3 minutes per location (total treatment time 24 minutes). In the nearby untreated region, a CH treatment was then performed with the same transducer, but with CH parameters: 7.7-µs pulses, 150-Hz PRF. A single row with 4 locations separated by 3 mm were treated for 5 minutes each (total treatment time 20 minutes).

Post-treatment, hyperechoic reflectors, presumably from residual histotripsy bubbles trapped in surrounding pus, could be observed outlining the CH and BH treatment zones (Fig 4b). A histological section of the same abscess is shown in Fig. 4c, where the BH and CH treatment zones have been labeled. Allowing for changes in the tissue as a result of tissue processing, the histological section correlates well with the bubble outlines.

Examination of the BH and CH lesions at higher magnification (Fig. 5) revealed histological features characteristic of histotripsy lesions. Both lesions contained homogenized cells with a lack of structural features found in untreated tissue. There was no discernable difference between BH and CH lesions. Dextran particles were observed in both the treated and untreated regions. In some treated regions, fragmented dextran particles could also be seen. Outside of the treatment zone, the pus consisted of a large number of viable and degenerating inflammatory cells, tissue fragments, bacteria and protein. One difference in the treated area in abscesses compared to organs is the presence of intact cells in the homogenized region.

**Figure 5:**
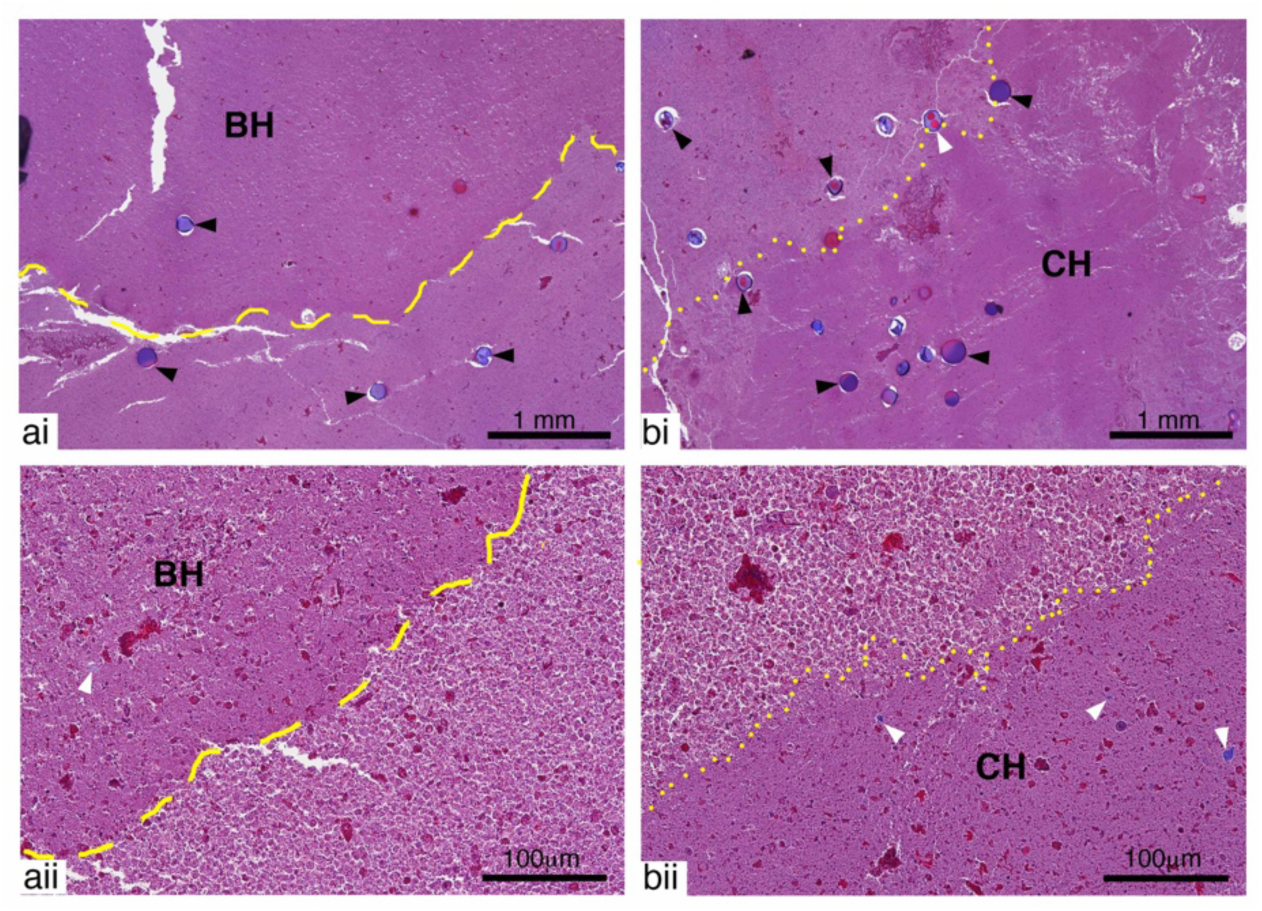
Representative images of Masson’s Trichrome stained sections showing a) BH and b) CH lesions at i) low and ii) high magnification. Both BH and CH lesions contained mostly homogenized pus lacking the structural features present in untreated tissue. Whole dextran particles (black arrow heads) were present in both treated and untreated areas. Fragmented dextran particles (white arrow heads) could also be observed in areas of the treatment region for both lesions. Outside of the treatment region intact and degenerating inflammatory cells are present.

Returning to the ultrasound images, the 2D images were reconstructed in a regular 3D grid using spatial tracking data to create a gray-scale volume; technique details can be found in the references (Leotta and Martin 2000). The borders of the abscess and the two treatment regions were manually outlined in a series of planes viewed in orthogonal directions, and the 2D outlines were used to construct a 3D surface for each object using the MeshLab software package (Cignoni, et al. 2008). In Fig. 6a, a slice though the reconstructed volume shows the two treatment regions in a unique “constant depth” plane, while in Fig. 6b, surface reconstructions created from outlines are shown for the abscess and the treatment regions. The white mesh represents the full abscess, the BH region is shown in red and the CH region in green. The BH treatment volume was 4.9 cc, while the CH treatment volume was 1.2 cc.

**Figure 6.**
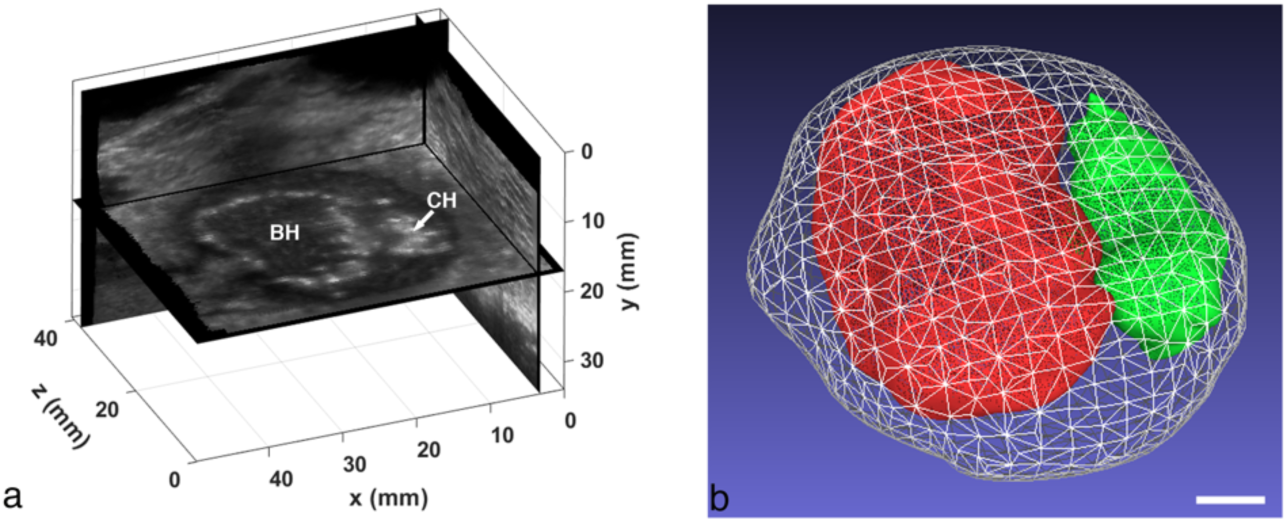
(a) Orthogonal views from a 3D volume reconstruction of the ex vivo abscess that was treated with BH and CH. The “constant depth” image plane showing the BH and CH regions is unique to the reconstruction. (b) Surface reconstructions of the full abscess (white mesh), BH treatment region (red) and CH region (green). The total abscess volume is 14.6 cc (BH volume: 4.9 cc, CH volume: 1.2 cc). Scale bar represents 1 cm.

The contents of both BH and CH regions were equally easy to aspirate with an 18-G needle following treatment, whereas the pre-treatment contents were substantially more viscous and more difficult to aspirate, indicating that liquefaction occurred following both treatments. Bacteria kill rates are discussed in the ‘bacteria reduction’ section below.

### In vivo studies

The pilot *in vivo* experiments were designed to (1) develop the workflow for *in situ* treatments, (2) evaluate treatment monitoring and post-treatment changes observed with ultrasound imaging, and (3) quantify bactericidal effects. Representative cases are provided.

The following case describes a treatment 3 weeks after inoculation, in which the animal was allowed to recover, eventually being euthanized 3 weeks after treatment (6 weeks after inoculation). Pre- and post-treatment ultrasound images obtained from the GE LOGIQ Book XP are shown in Fig. 7. The relatively uniform core is surrounded by a dark vascularized capsule, which itself is outlined by a brighter echogenic rim.

**Figure 7.**
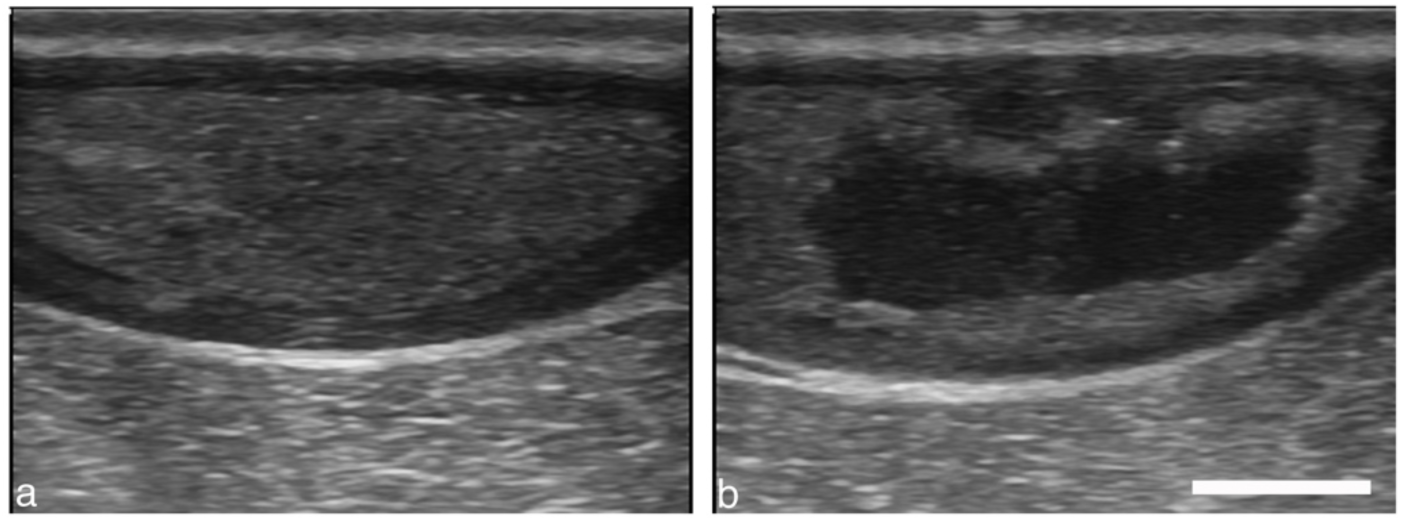
Intramuscular abscess (a) before and (b) after a 20-min CH treatment with 50 cycle bursts at 300 Hz PRF. The abscess size is 4.8 cm x 1.6 cm in this plane. The treatment zone was scanned manually by observing and maintaining the cavitation cloud within the abscess. Scale bar represents 1 cm.

The intramuscular injection grew into what was deemed a mature abscess at week 3 when CH treatment was performed with the 1.5-MHz, F# 0.77 transducer. After 3 minutes of treatment, the probe was manually moved 1-5 mm without pausing as treatment continued for a total of 20 min. Even with limited imaging quality, a cavitation cloud could be seen during treatment, and post-treatment imaging of the abscess revealed a treatment zone covering about half the abscess. The aspirate was less viscous post-treatment, indicating that histotripsy liquefied the viscous contents.

Because of the difficulty in associating treatment bioeffects with histology that was performed weeks after treatment, later studies focused on euthanizing the animal immediately after treatment and taking tissue sections for histological analysis so that more direct comparisons could be made.

A BH-treated abscess is shown in Fig. 8. This subcutaneous abscess was located in the lower abdomen of the pig. Figure 8a shows a B-mode image before treatment. The 3D surface reconstructions of the abscess and treated region are shown in Fig. 8c. The outer white mesh corresponds to the abscess with capsule, and measured 125 *x* 71 *x* 43 mm. The yellow mesh is the abscess core (109 *x* 61 *x* 35 mm) and the red mesh is the treated region (101 *x* 47 *x* 29 mm). In this case, respiratory motion resulted in significant misregistration of the images acquired with the 3D tracker. Therefore, the 3D reconstruction was approximated by compiling the image series into a 3D volume assuming a constant speed linear scan, with the total scan length provided by the 3D tracker. Figure 8d illustrates a SWE image of the treated abscess. The box spans both the treated and untreated regions. The untreated region has a mean elastic modulus of 4.5 kPa, while the treated area has a mean elastic modulus of 1.2 kPa. The treated region barely supports a shearwave indicating it is likely liquified, which was further confirmed upon aspiration.

**Figure 8.**
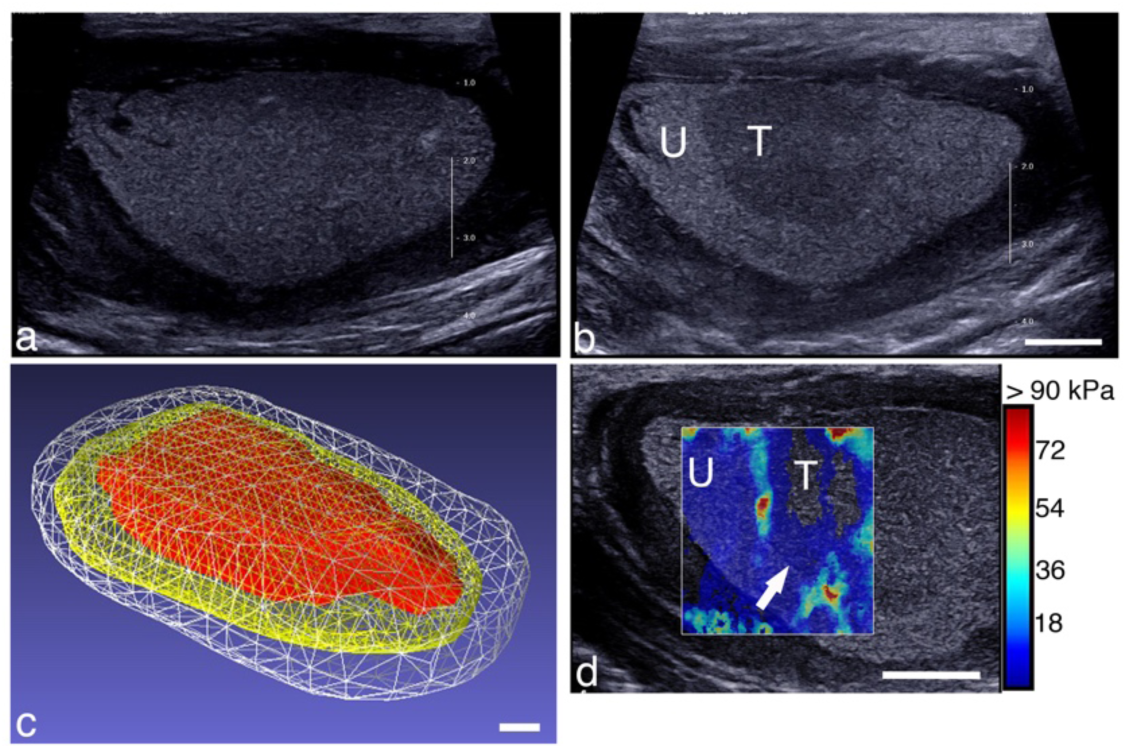
B-mode image of a subcutaneous abscess (a) before and (b) after BH treatment. The treated area (labeled ‘T’) looks qualitatively different than the untreated region (labeled ‘U’). The treatment zone was scanned with the robotic arm with 3-mm spacing between focal spots, 1 min per location, by observing and maintaining the cavitation cloud within the abscess. (c) Reconstructed volume of the abscess and treated region. The abscess volume with capsule (white) was 213.7 cc, while the core volume (yellow) was 127.7 cc, and the treatment zone (red) was 61.2 cc. (d) Shearwave elastography image overlaying a portion of the treated and untreated regions. The untreated region’s stiffness was 4.5 kPa, while the treated region measured 1.2 kPa. Scale bars represent 1 cm.

Another example of a CH-treated abscess is shown in Fig. 9. This particular abscess had a very thick capsule distally. Surface reconstructions of the abscess and treated region (Fig. 9c) show that the abscess size was 61 × 38 × 29 mm with capsule, while the core was 60 × 32 × 18 mm. The treatment zone was 23 × 20 × 14 mm. This 3D reconstruction was approximated by compiling images from a freehand sweep into a 3D volume assuming a constant speed linear scan, with the total scan length based on a pre-treatment tracked 3D scan.

**Figure 9.**
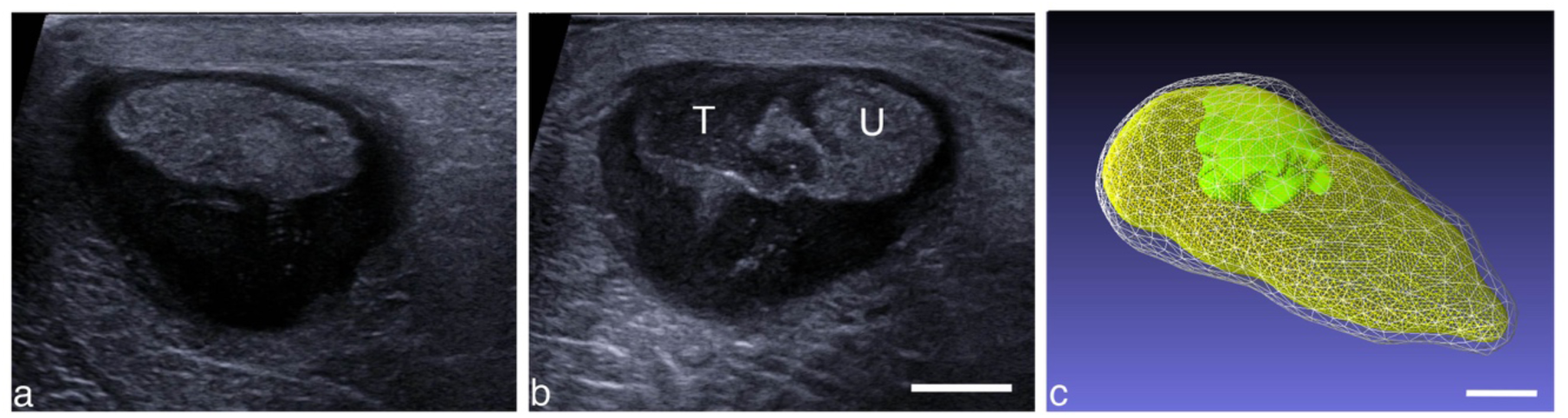
Intramuscular abscess (a) before and (b) after a CH treatment. The abscess size is 61 × 38 x 29 mm. The treatment zone was located manually by observing and maintaining the cavitation cloud within the abscess. U (T) corresponds to untreated (treated) regions. (c) Reconstructed volume of the abscess and treated region. The abscess volume with capsule (white) was 26.7 cc, while the core volume (yellow) was 13.1 cc, and the treatment zone (green) was 2.1 cc, scanned using the robotic arm with 3-mm spacing and 3 min per focal spot. Scale bars represent 1 cm.

A SWE image across the zones of treated and treated abscess is shown in Figure 10b. The untreated portion of the abscess was heterogenous in elasticity with a mean elastic modulus of 11.2 ± 2.7 kPa, while within the treated area the stiffness was markedly reduced, 2.0 ± 0.6 kPa.

**Figure 10.**
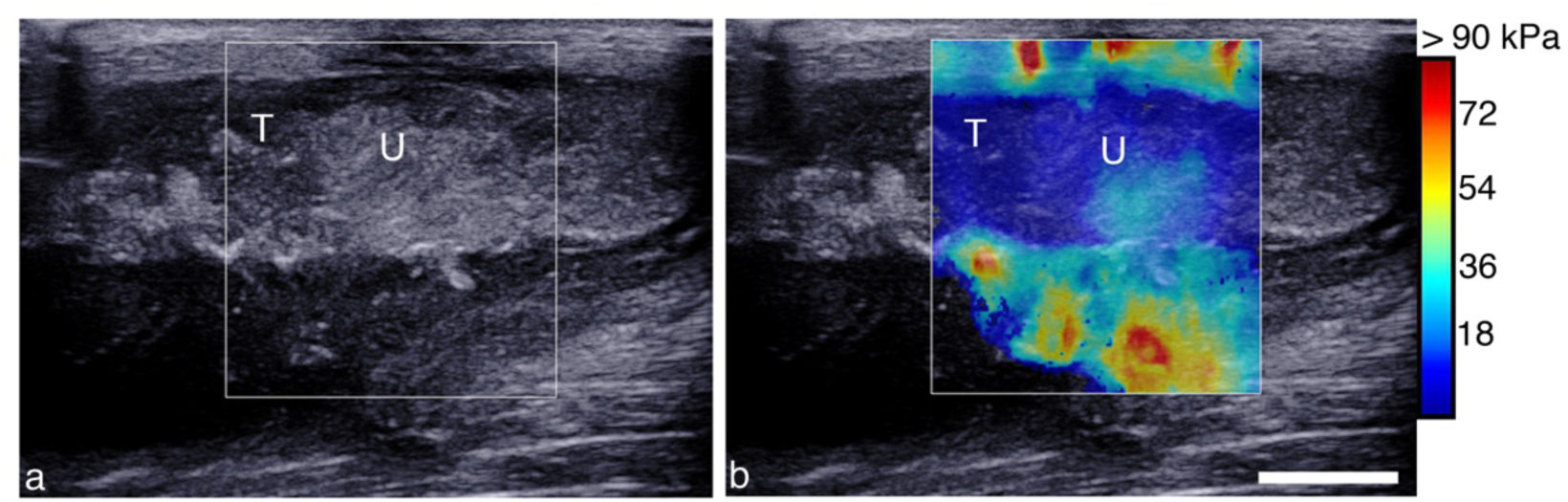
Post treatment B-mode (a) and SWE (b) image across the boundary of CH showing changes in echogenicity and reduction in elasticity. (a) B-mode image shows an echogenic border demarking the untreated region (U) from the central treated part of the abscess (T). (b) SWE image measuring 2.0 ± 0.6 kPa in the treated region (T) and 11.2 ± 02.7 kPa in the untreated region (U).

The fibrous capsule contains vessels (see *e*.*g*. Fig. 4c and Figs. 11-12). Application of either CH or BH treatment near the capsule-cavity border resulted in damage to the inner capsule vasculature (Fig. 11b,c), resulting in varying degrees of hemorrhage and thrombosis. In many cases, bleeding into the abscess cavity was observed in the post-treatment aspirate, as indicated by a change in the hue of the pus withdrawn. Pre-treatment pus was often a pale yellowish-green; the post-treatment pus was often tinged with a pale pink hue most likely due to bleeding from damage to the inner portion of the capsule. In the samples analyzed, damage never extended completely through the capsule. Fragments of the internal surface of the collagenous capsule were often seen within the cavity of the abscess (Figs. 11bii,cii).

**Figure 11:**
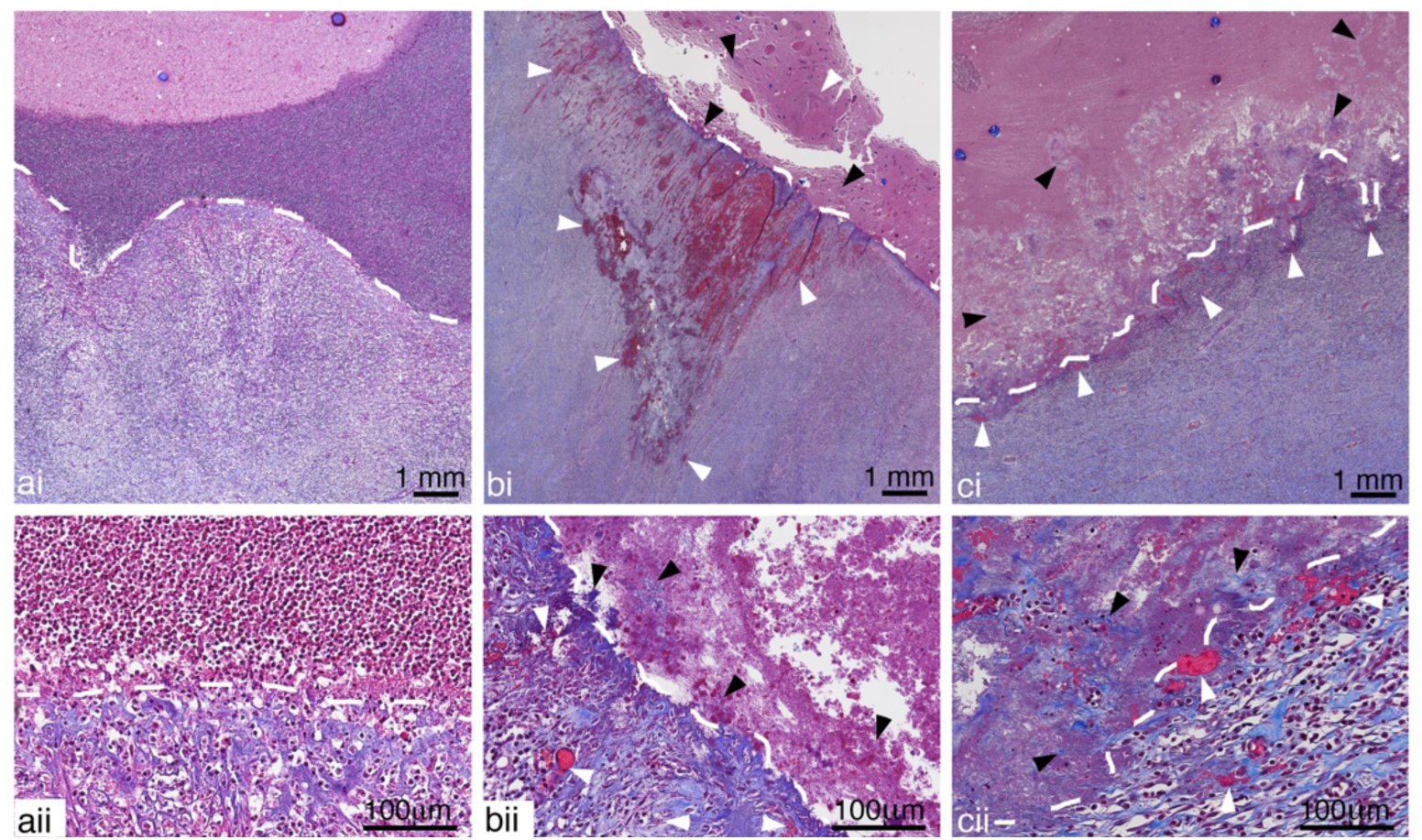
Representative images of Masson’s Trichrome stained sections showing a) untreated, b) CH and c) BH treated abscesses close to the inner portion of the capsule (white dashed lines) at i) low and ii) high magnification. A well-defined border in the untreated abscess shows organized collagen fibers and intact vessels within the capsule and a dense collection of inflammatory cells in the adjacent regions inside the abscess cavity. Vessel damage (white arrow heads), with blood leakage and thrombosis, can be observed in the fibrous capsules of both BH and CH treated abscesses. The cells in the abscess cavity close to the capsule are homogenized and fragmented collagen fibers (black arrow head) torn away from capsule can also be seen. The damage did not extend to the outer regions of the capsule.

The cavity of infected abscesses contains pus consisting of inflammatory cells, degenerating cells, tissue fragments (e.g. muscle, fat, connective tissue), proteinaceous material and bacteria (Wang, et al. 2020). Within the cavity, and similar to the *ex vivo* histotripsy lesions, there were some larger regions of homogenized cells with a generally well-defined border for both the CH and BH lesions (Figs. 12b,c). Unlike lesions generated in soft tissues, the homogenized regions also contained a number of intact cells and there were smaller collections of homogenized pus observed throughout the abscess cavity, outside of the main lesion (Figs. 12b,c). Within the treated zone there was presence of fragmented tissue (smaller fragments compared to regions outside of the homogenized lesions) and collagen depending on what tissue pieces were present in the abscess prior to treatment (Fig. 12b).

**Figure 12:**
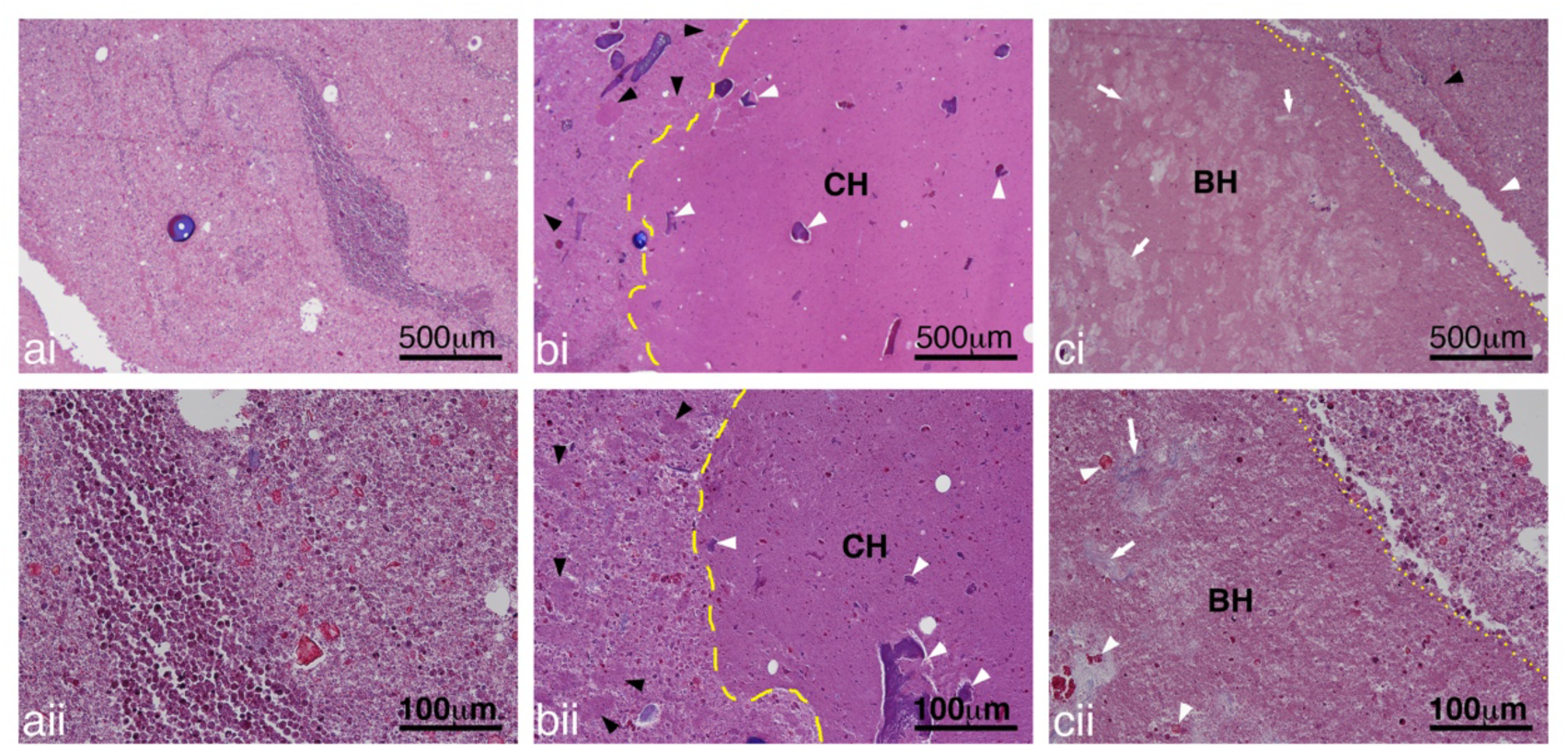
Representative images of Masson’s Trichrome stained sections showing a) untreated, b) CH and c) BH treated abscesses at i) low and ii) high magnification. The pus in the untreated abscess consisted of a mix of inflammatory cells, degrading inflammatory cells, tissue pieces and proteinaceous material. At higher magnification cellular structure can still be observed even for the degrading inflammatory cells. In the treated abscesses, there were some larger regions of homogenized cells with a generally well-defined border for both the CH (yellow dashed line) and BH (yellow dotted lines). Smaller collections of homogenized pus (black arrowheads) was present outside of the main lesion. Within the lesions there was presence of damaged tissue (white arrowheads) and collagen (white arrows).

### Bacterial reduction

In total, 15 bacterial reduction treatments were performed. However, in 4 cases we were unable to aspirate liquid pre-treatment using the 18-G needle. In 14/15 cases, aspirate was removed post-treatment, indicating liquefaction of the contents. For quantification, pre- and post-treatment aspirate pairs were successfully obtained and the bacteria quantified in 3 CH treatments (all *in vivo*), 4 BH treatments (one *ex vivo*, the rest *in vivo*) and one CH + BH treatment (*ex vivo)*. Data were quantified in terms of the relative number of microbes eliminated by treatment. It is defined as Log Reduction = log_10_(N_0_/N), where N_0_ is the number of colony forming units (cfu) before treatment, and N is the number of cfu after treatment.

Figure 13 shows the log reduction in bacterial load for each histotripsy regime. The specific reductions for CH were Log [0.7, 0.8, 1.1]; BH reductions were Log [1.2, -0.9, -0.4, 0.1]. Although not shown here, a combination of CH followed by BH treatment that essentially doubled the total treatment time, led to a Log kill of 3.3.

**Figure 13.**
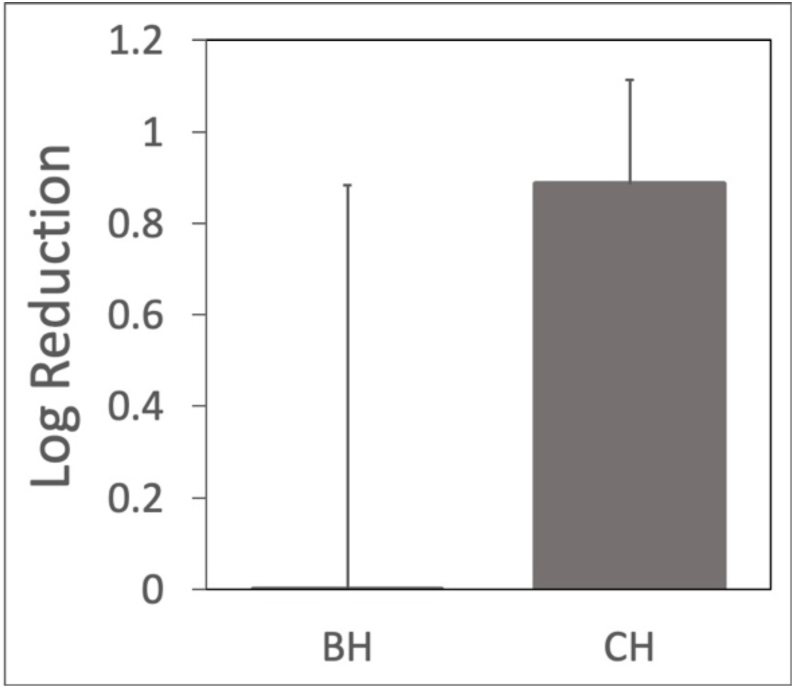
Log reduction in bacteria viability. CH consistently demonstrated bactericidal effects, while BH generated mixed results. A single application of CH and BH that doubled the treatment time resulted in a 3.3 Log kill (not shown).

### Transmission Electron Microscopy (TEM)

Figure 14 shows representative TEM images of abscessal contents sampled before and after treatment. Untreated abscessal contents contained visibly intact gram negative bacteria, intact and degenerating eukaryotic cells evident from the sea of non-constrained organelles. Samples taken from the treatment zone contained a homogenized cellular debris. The cytoplasmic cell membranes were absent and cellular components were mostly indistinguishable. Ghosts of organelles could be observed in the form of membranous structures amongst a sea of cellular debris. There were a few cells that were not completely homogenized, although they were scarce compared to homogenized cells.

**Figure 14.**
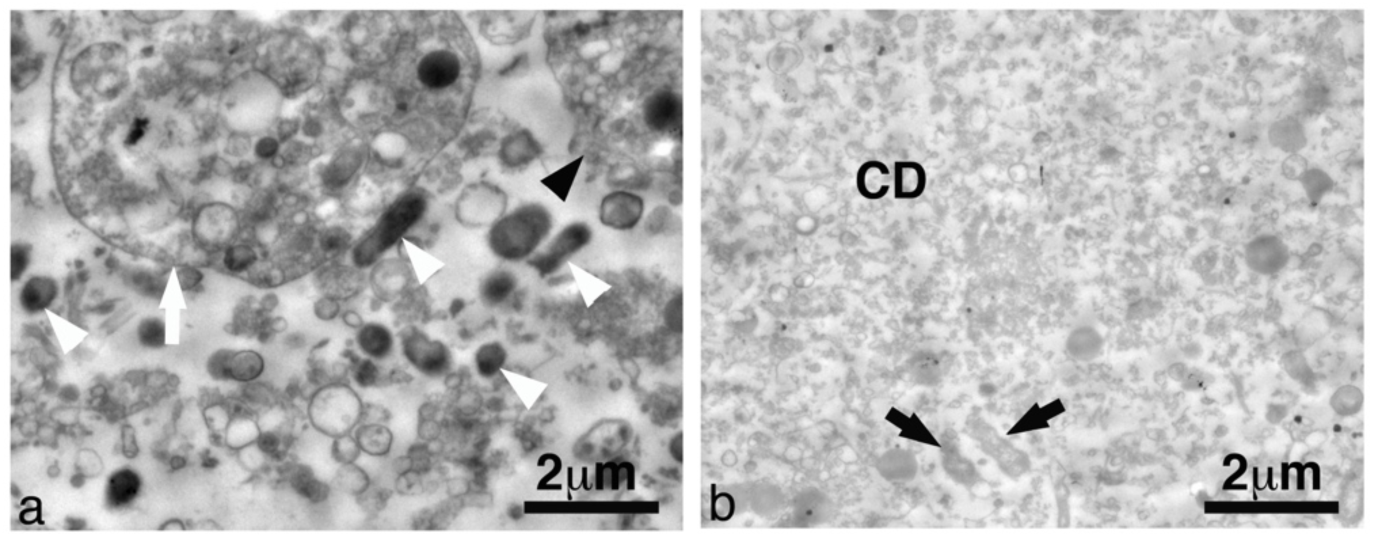
Representative transmission electron microscopy images from abscessal contents taken (a) before and (b) after histotripsy treatment. This particular abscess was treated with CH. Untreated abscessal samples contained intact cells (black arrowhead), degenerating cells (white arrow), intact bacteria (white arrowhead) and un-constrained organanelles. Treated abscessal samples contained a sea of mostly homogenized cellular debris (CD) with a few damaged bacteria (black arrow).

## Discussion

A pilot study was conducted to evaluate the application of focused ultrasound to abscesses grown in a novel porcine animal model. *Ex vivo* and *in vivo* studies were performed to assess the effects of two types of histotripsy treatments: CH and BH. Acoustic and histopathological evaluation of treated abscesses was performed in addition to an evaluation of the bactericidal effect. Several observations are noted here.

### Identifying mature abscesses

The injection of bacteria and dextran initially resulted in the development of a phlegmon that then evolved into an organized mature abscess over 2-4 weeks. B-mode ultrasound allowed us to identify key structural characteristics, namely a well-defined hypoechoic capsule surrounding a core with mixed echogenic material, which were used as an indicator of maturity (Lu, et al. 2019). Doppler imaging was used to further evaluate abscess maturity, with mature abscesses having rich vasculature within the hypoechoic capsule, and an avascular core. This vascularized capsule should correspond to the enhancing rim used as CT criteria to suggest an abscess. In general, the pig abscesses were more echogenic than often observed in humans, although echogenicity of human abscesses can be variable (Taylor, et al. 1978, Sarrazin and Wilson 1996, Esteban, et al. 2003, Lubbert, et al. 2014). One contributing factor may be the addition of dextran particles (∼150 µm) used to localize the abscess, which likely increases scattering and makes the abscess appear brighter. Dextran particles may also act as cavitation nuclei and enhance cavitation activity. However, bacteria create gas as a byproduct of their digestion; and abscesses are not degassed. Also, treatment pressure levels are high enough to generate cavitation in even degassed liquids. For these reasons, we hypothesize that the presence of dextran does not enhance cavitation activity.

### Histological features

The pig abscesses generated in this model have salient features observed in human abscesses (Kumar, et al. 2015). The mature abscesses have a well-defined fibrous connective tissue capsule with a cavity filled with aggregations of neutrophils within a field of proteinaceous and degenerating inflammatory cells, bacteria, and in some cases tissue pieces. Unlike human abscesses, these abscesses also contain dextran. The purpose of the dextran here was to elicit a persistent inflammatory response to aid in the reliable generation of an abscess (Wang, et al. 2020). In some cases dextran fragments were found in the treated abscesses. Fragmentation of the dextran likely resulted from the large shear forces generated by the collapsing of the cavitation bubbles.

Dextran aside, histological evaluation of both CH and BH treatment regions revealed histological features characteristic of histotripsy lesions (Maxwell 2012, Wang, et al. 2013, Khokhlova, et al. 2014, Khokhlova, et al. 2016, Wang, et al. 2018). Both lesions contained a large portion of homogenized cells with a lack of structural features found in untreated abscesses. However, unlike lesions generated in soft tissues, the homogenized regions also contained a number of intact cells and there were smaller collections of homogenized pus observed throughout the abscess cavity, outside of the main lesion. We hypothesize that this is due to the mixing that happens as a result of the treatment. There was no discernable difference between BH and CH lesions. At the ultrastructural level, electron microscopy supported the histology observations, but revealed damage to the cells and bacteria at an ultrastructural level.

### Treatment observations

Although the characteristic features of histotripsy lesions are present, there are several differences in the lesions generated in ‘liquid’ abscesses compared with those generated in ‘solid’ tissues. Unlike lesions generated in soft tissues, the homogenized regions also contained small islands of intact pus and tissue, and conversely, collections of homogenized pus could also be observed throughout the abscess cavity, outside of the main lesion (*e*.*g*., Fig. 13b). Both of these observations are likely due to the macroscopic streaming which circulates cells and other particles in the liquid pus through the focus (Brayman, et al. 2018). It is also possible that post-treatment aspiration and post euthanasia tissue processing could cause some mixing of the treated and non-treated pus.

Although fibrous tissues are accepted to be more resilient to histotripsy treatment (Vlaisavljevich, et al. 2014), there was evidence that fibrous tissue was damaged by our treatment (see, *e*.*g*., Fig 12b,c). This suggests that not only can histotripsy be used to liquify pus for fine needle aspiration, it could potentially break up septal walls that separate multiple compartments in loculated abscesses. Clinically, septations sometimes require separate drains into each compartment. Histotripsy may thus be useful as an adjunctive therapy to break up septations, allowing for a single drain and reducing complications.

One concern with histotripsy is that mistargeting onto the capsule itself might rupture it and lead to the infection spreading. Ideally, macroscopic streaming obviates the need to treat near the capsule cavity border as surrounding liquid pus becomes entrained in the streaming and moves through the focus. No attempts were made to evaluate microstreaming treatment at the capsule, but treatment near the capsule is evidenced in Figs. 4, 8, and 9. Histological analyses of the capsule revealed some capsular damage (Fig.11). Although regions of vessel damage and some superficial collagen damage was observed, the capsule remained intact and there was no leakage of the abscessal contents outside of the cavity that could be seen. Unlike catheter drainage, which pierces the entire capsule, histotripsy did not damage the outermost regions of the capsule.

Turning to the SWE images, we found the treated regions barely supported a shearwave indicating near or complete liquefication. The liquid state of the treated regions was confirmed upon aspiration. The pre-treated lesions appeared heterogenous in stiffness, which was mostly homogenized to a 1-3 kPa after treatment. For reference, normal liver tissue is ∼5-6 kPa in stiffness. The surrounding muscle was observed to be 7-10 kPa depending on the orientation to the fibers. In Fig. 8c, the vertical line between the treated and untreated regions is most likely an artifact associated with the shearwave reflection between the interface of these regions (I.e. stiffer and liquid regions respectively). Conventional SWE processing assumes a homogenous medium where the shearwave is propagating in a single direction and the shearwave speed is then directly proportional to the shearwave modulus. The presence of a strong reflected wave propagating in the opposite direction breaks this assumption. Future work is planned to study the effect of liquid-solid boundaries on SWE images.

### Liquefaction

All post-treatment aspirates were qualitatively less viscous than the pre-treatment aspirates, as determined by the ease/speed of aspiration. In several abscesses, aspiration was only possible after treatment. This qualitative finding suggests a possible treatment in which the pus is liquefied and removed by a fine needle, rather than requiring a relatively larger drain. Additional evidence for liquefaction comes from the B-mode images, where treated regions were less echogenic, suggesting homogenization of the contents. The average reduction in B-mode pixel intensity from 9 separate treatment regions was 34 ± 20 pixels.

### Log reduction in bacterial load

Generally, BH treatments resulted in liquefaction, but were not as reliable in bacterial inactivation. We attribute this partly to treatment dose, as described above. Ultrasound inactivation of bacteria has a strong dose dependence (Brayman, et al. 2017, Brayman, et al. 2018). The microstreaming would change the effective treatment dose as pus is moved through the focus. Over similar treatment times, CH recirculates material at a faster rate than BH. CH was successful in producing a mean log kill of 0.89 ± 0.23 (SD). This should be improved by longer treatments, or the combined treatment of BH and CH. Indeed, the one abscess that was serially treated with the two types of histotripsy yielded a 3.3 log kill (99.9% reduction in viable bacteria).

One final observation worth noting is of the abscess that was left to progress for 3 weeks after treatment. In a different imaging plane from the same treatment, a tract is seen extending from the abscess to near the skin surface (Fig. 15a). This distinct track may be associated with the needle tract formed by the post-treatment aspiration. Interestingly, there is a corresponding track observed in the histology, obtained after the animal was euthanized 3 weeks after treatment. We speculate that these tracks may be related, and if so, this would be direct evidence that bacteria can move up the needle track. An alternative explanation is that the track observed in histology may have come from a fistula developed naturally, as the abscess was allowed to evolve for 3 additional weeks post-treatment. Note too that by the time the animal was euthanized, the abscess itself was much smaller, indicated by the small red volume of material seen in Fig. 15b. Unfortunately, the track apparently provided an outlet for pus expulsion through the skin in the weeks following treatment, so it was impossible to know how much shrinkage was due to spontaneous resolution by resorption *vs*. how much was due to pus expulsion. Parenthetically, if the track was due to needle aspiration, it has important implications for the current standard of care, which uses large bore drains during percutaneous drainage to remove pus and bacteria. Indeed, secondary infections, a common sequela of drainage, are most likely related to bacteria moving along the drain tract. Histotripsy-induced liquefaction may allow the clinician to use small-gauge needles for substantive aspiration, reducing the potential for secondary infections.

**Figure 15.**
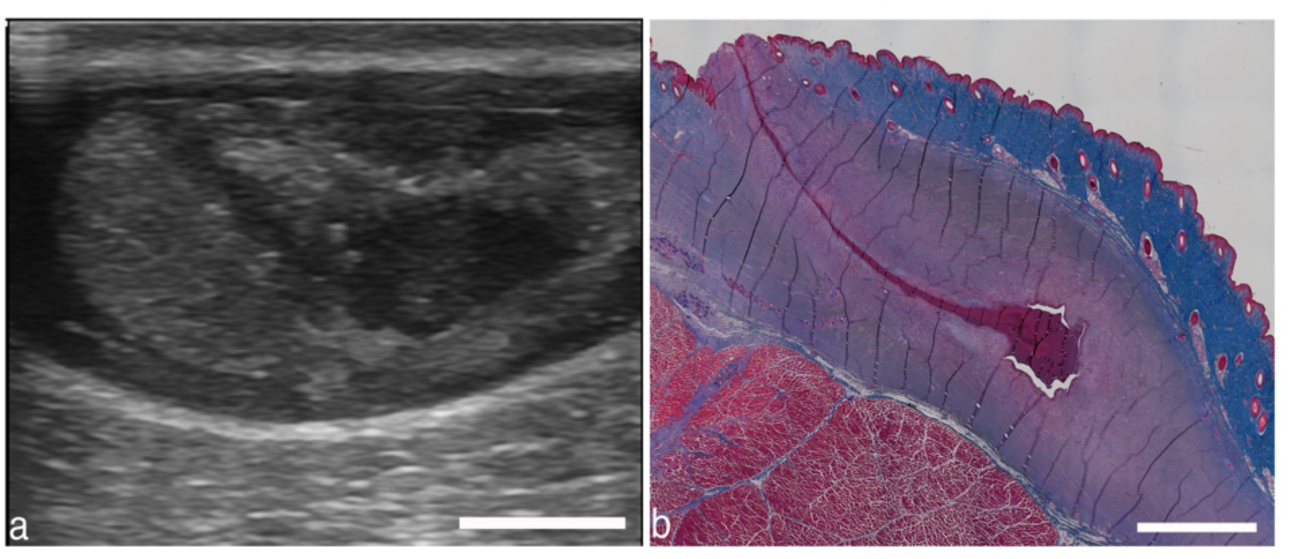
(a) Different image plane from abscess in Fig. 8. A darkened region extending from the abscess upwards at an angle may be due to the needle used to aspirate contents before and after treatment. (b) Masson’s trichrome stained section from the same abscess from 3 weeks after treatment shows a similar track. The track may be from the needle aspiration, indicating a potential track for bacteria to escape the abscess. Alternatively, the track may be a naturally formed fistula. Scale bars represent 1 cm.

### Limitations

The goal of this pilot study was to evaluate the use of focused ultrasound to treat abscesses. Despite encouraging results, there were a number of limitations that need to be addressed in future studies. The abscesses were only partially treated so that ultrasound imaging could be used to assess gross changes in the treatment zone. Given the low viscosity of the pus in some of the abscesses, it is likely that mixing of the treated and untreated pus occurred during and, due to necessary manipulations, also after treatment, leading to measurement error in the post-treatment colony counts. Indeed, histological evaluation supports this with islands of intact pus found within the lesion. Despite only partially treating the abscesses and evidence of mixing, it is encouraging that we observed a mean log kill of 0.89 ± 0.23 (SD) for the CH treated abscesses. In the majority of the abscesses, only one sample was taken pre- and post-treatment. This was to minimize the disruption of the treated regions for the histopathological evaluation.

## Conclusion

A histotripsy pilot study was successfully carried out on abscesses in a novel porcine model. Furthermore, novel approaches to imaging were applied to provide assessment of the abscess and real time treatment guidance. Abscesses represent a class of infected fluid collections which differ from solid tissues, as streaming, mixing, and heating rates vary greatly. Abscesses matured over 2-6 weeks, reaching sizes up to 12.5 *x* 7.1 *x* 4.3 cm. Boiling and Cavitation histotripsy effectively liquefied viscous abscess contents. Cavitation histotripsy resulted in higher bacterial kill rates, most likely due to higher treatment dose. Histotripsy is a potential adjunctive therapy to percutaneous drainage by breaking septations and reducing the need for multiple drains. Histotripsy-induced liquefaction of viscous pus may allow simple fine needle aspiration that obviates the need for long-term catheter drainage or surgical evacuation, reducing complication rates, morbidity, hospital length of stay and improving quality of life for patients with clinically significant abscesses.

## Acknowledgements

Work supported in part by NIH grants 5R01EB019365, R01GM122859 and K01 DK104854, and RSF grant 20-12-00145. The authors wish to thank Ivan Pelivanov and Matthew O’Donnell for discussions on SWE imaging.

